# Lateral pressure profile in bent lipid membranes and mechanosensitive channels gating

**DOI:** 10.1101/062463

**Authors:** Anna A. Drozdova, Sergei I. Mukhin

## Abstract

This review describes the analytical calculation of lateral pressure profile in the hydrophobic part of the lipid bilayer with finite curvature based on previously developed microscopic model for lipid hydrocarbon chains. According to this theory the energy per unit chain is represented as energy of flexible string (Euler’s elastic beam of finite thickness) and interaction between chains is considered as an entropic repulsion. This microscopic theory allows to obtain expression for lateral pressure distribution in bent bilayer if treating a bending as a small deviation from the flat membrane conformation and using perturbation theory. Because lateral pressure distribution is related to elastic properties of lipid bilayer then the first moment of lateral pressure and the expression for bending modulus may be derived from this theoretical model. Finally one can estimate the energy difference between two various conformational states of mechanosensitive channel embedded into the bilayer with pressure profile П_*t*_(*z*).

---

The relevance of the present study is supported by the fact that biological membranes constitute complex supramolecular structures that surround all living cells and form them in closed organelles. The basic structural element of the membrane is lipid bilayer, whose mechanical and thermodynamical properties are believed to play important role in the functioning of organized matter.

The lipids have one polar (hydrophilic) “head” and two non-polar (hydrophobic) hydrocarbon “tails” (chains). There is a great variety of membrane constituents and lipid characteristics (head group type, chain length, etc). Our analytical theory of lipid membranes, based on the microscopic model, was developed in [1,2]. In this model each lipid chain is modeled as a flexible string of finite thickness (incompressible area per chain) with given bending rigidity (the case of persistence length comparable with the chain full length). The ensemble of chains is investigated within a mean-field approximation with respect to the strings collisions. According to this approximation a steric (entropic) repulsion between the lipid tails is represented with an effective lateral potential acting on each string in each monolayer. The negative lateral tension keeping all the strings (i.e. lipids) together comes from hydrophobic-hydrophilic interface in the head-groups area of a membrane and is treated as an input parameter of the model. So, the energy functional for a lipid chain can be written analytically and then partition function and free energy of the chain can be estimated. The strength of the effective lateral potential acting on each string, that minimizes the free energy, is found then in a self-consistent (mean-field) manner. The analytical expression of the free energy allows to obtain different macroscopic characteristics of lipid bilayer, particularly, the lateral pressure profile that proves to be very important feature of lipid membrane because it is related to the elastic properties of lipid bilayer [3] and also the functionality of mechanosensitive ion channels [4, 5, 6].

Mechanosensivity is a general sensory mechanism found in living organisms. It’s known that the functionality of many membrane proteins, such as MscS (mechanosensitive ion channels of small conductance), and the most studied bacterial membrane protein MscL (mechanosensitive ion channel of large conductance), is sensitive to the lipid environment [7, 8, 9] and form mechanically gated channels when they embedded in a lipid bilayer. In each organism mechanosensitive channels (MSCs) perform the certain functions. For people MSCs are responsible for the regulation of bloodpressure, take hearing and vestibular function, and also play a role of touch receptors [10].

Because mechanosensitive channels are inserted in a lipid bilayer, they are sensitive to its deformations [11]. MSCs respond to mechanical stress by changing of the channel shape between the closed and open state. The change in external dimensions between the closed and open states associated with the opening of a pore in MSC. [9], [12].

Lateral pressure profile have been suggested to play a significant role in activation of mechanosensitive channel through changes in its conformational state [13, 14, 15]. The problem is that experimental studies of the lateral pressure are rare and indirect [16]. So in this research we focus on analytical derivation of expression for the lateral pressure and comparison our results with computational calculations of other colleagues.

Our microscopic theory is quite general, but for comparison of the analytically obtained lateral pressure with experimental data we use the reference typical microscopic parameters of e.g. DPPC lipids: the monolayer thickness L_0_=1,5 *nm*, incompressible area per chain A_0_=0,2 *nm*^2^. The flexural rigidity of a lipid tail can be evaluated then from the Flory’s polymer theory: *K_f_*≈*k_B_T*_0_*L*_0_/3, where temperature is *T_0_*=300 *K*. To compare with experiment the analytically obtained values for the first moment of the lateral pressure κ*_m_J_s_* (here κ*_m_* is the monolayer bending modulus and *J_s_* is the monolayer spontaneous curvature), and bending modulus of bilayers κ*_b_* we calculate parameters of lipid molecules for different fully saturated lipids. Also we calculated a work associated with embedding the mechanosensitive channel in its different conformational states into various lipid bilayers. Obtained expression depends on inter-facial surface tension, the channel area change in region between hydrophobic interior and the aqueous surroundings and also the bending modulus and spontaneous bending moment.

## Methods. Microscopic model of the lipid bilayer

Microscopic model of lipid membrane as an ensemble of flexible strings was investigated in [1,2]. Below we describe the basic statements of our theoretical model.

Hydrocarbon lipid chain is considered as Euler’s elastic beam of finite thickness (see fig. 1), and its energy functional is written in the following form: 
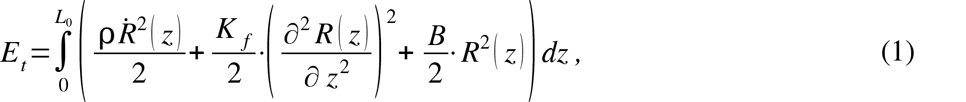

here ***R*** (*z*) is the deviation of the beam from the straight line at each level *z*, *L*_0_ - the monolayer thickness.

**Figure 1.**
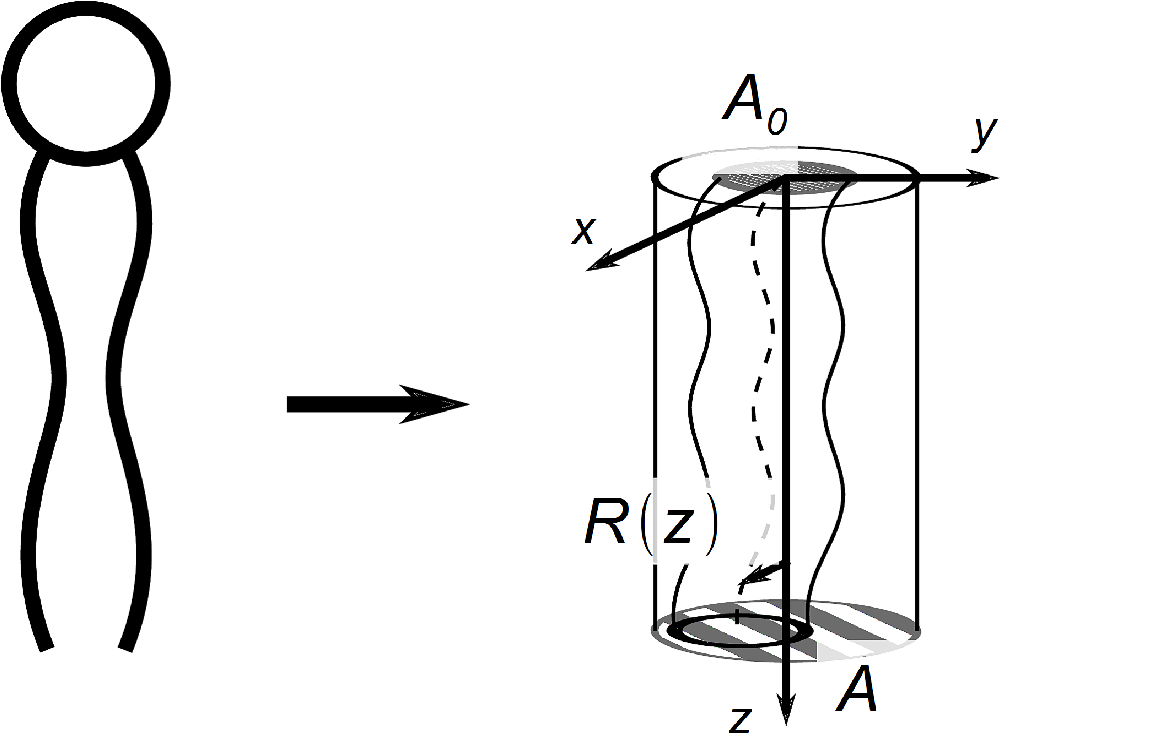
Hydrocarbon tails of lipid as flexible strings. *R (z)* - the deviation of the string from the cylinder axis, *A*_0_ is incompressible area per chain. Because the lipid strings makes the bending fluctuations its real area is greater than incompressible one: *A=A*_0_ *a*, where *a* is dimensionless area per lipid chain.

In formula (1) the first term is the kinetic energy of the chain, the second term is the bending energy of the chain, the third term is the potential representing entropic repulsion with the neighboring chains; ρ is the linear density of the chain.

Energy functional of the chain conformations (1) can be rewritten in the operator form, provided there are certain boundary conditions at the chain ends [1, 2]:

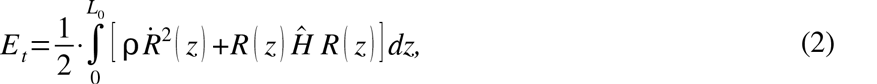

where the conformational energy operator is determined by expression: 
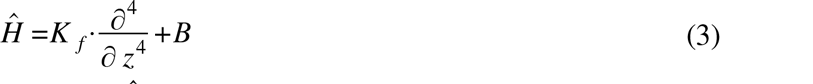

We can solve the operator equation *ĤR_n_=E_n_·R_n_* and find the eigenfunctions and eigen values of *Ĥ*. Then we expand the chain deviation ***R***(*z*) in these eigenfunctions: 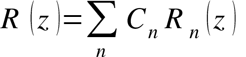. Using the orthonormality condition 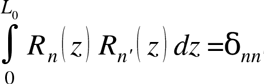, we can write the energy functional in the amplitudes {*C_n_*} representation of the expansion over the eigenfunctions:

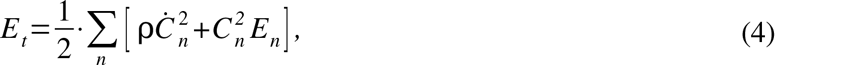

where *E_n_* is the n-th eigenvalue:

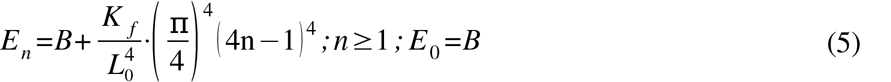
of the n-th eigrnfunction *R_n_*(*z*):

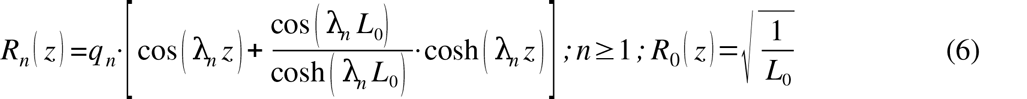

In last formula were used the following notations: 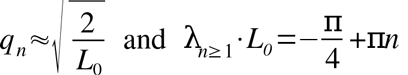.

Knowing the energy functional (1) allows one to calculate the partition function consisted in the product of two equal components, 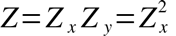:

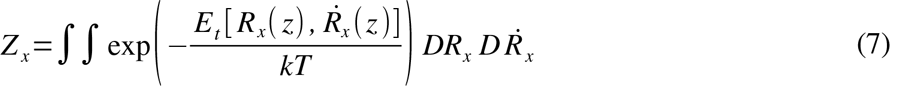

The partition function in the amplitudes {*C_n_*} representation takes the form:

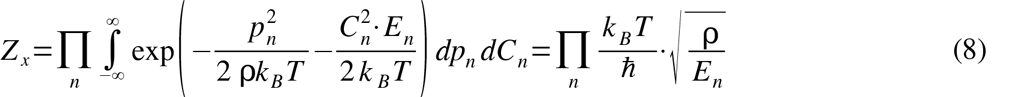

Hence, using the free energy formula:

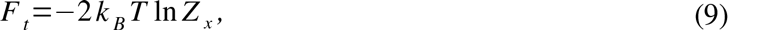

where *F_t_* is the free energy of the tails, and then substituting (8) into (9) one can obtain: 
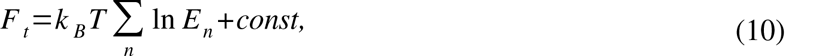

Using formula (10) we find the lateral pressure inside the hydrophobic part of the bilayer *P_t_*:

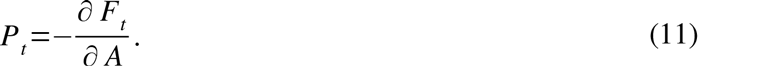

The self-assembly condition (i.e. zero total lateral tension of the self-assembled membrane) is represented in the form:

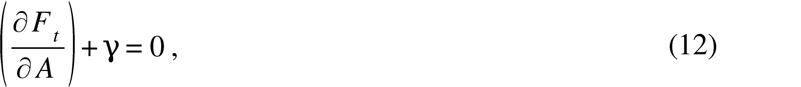

where γ is the surface tension at the interface between the polar and hydrophobic regions of the membrane [1]. The physical meaning of this value is the surface tension at the lipid-water interface (situated between hydrophobic tails and hydrophilic heads): γ=30-100 *dyn/cm* for various lipids. So the equilibrium condition (12) simply states that repulsion between chains should be balanced by the surface tension γ.

Lateral pressure is expressed in the following form [1]:

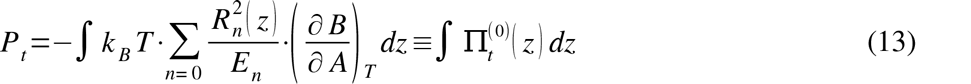

Here the lateral pressure profile distribution 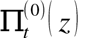 in the flat bilayer is introduced by the formula:

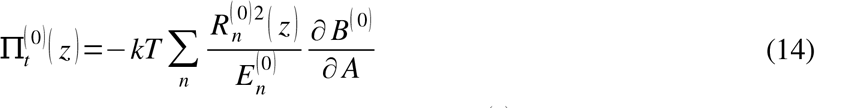
where the entropic repulsion coefficient in a flat membrane *B*^(0)^ and the area per lipid chain *A* can be found from self-consistency equation. Below we describe some details of this derivation.

The partial derivative of the free energy *F* with respect to *B* is:

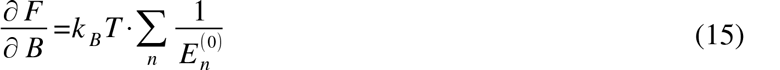

On the other hand, from the initial expression for the energy functional of the chain (1) the formula for partial derivative is expressed as follows:

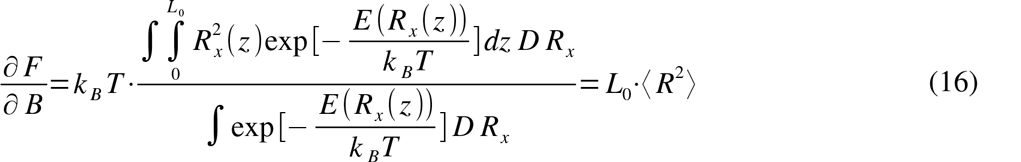

Here ⟨*R*^2^⟩ is the average of square deviation. It can be written in the form [1]: 
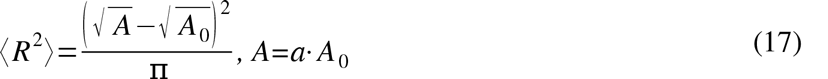

where *A*_0_ is incompressible area per lipid chain and *A* is the real area occupied by the chain.

Equating the right sides of the Eqs. (15) and (16) one gets the self-consistency equation: 
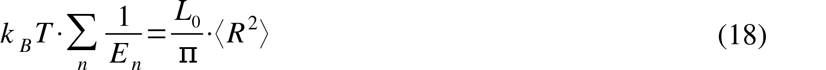

One can substitute the expression for the eigen values (5) into last formula (18) and obtain the self-consistency equation in dimensionless form [17]: 
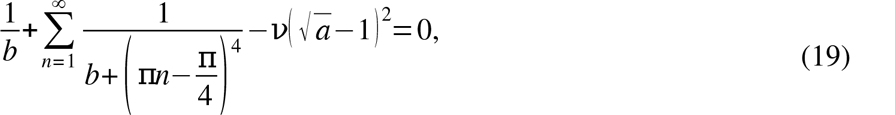

This sum might be rewritten as an integral 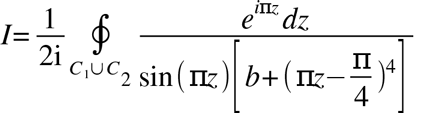. Evaluating this integral using residue theory one can obtain [18]: 
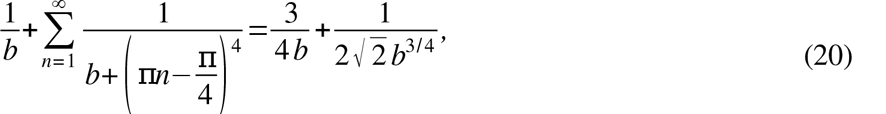
where *v* and *b* are dimensionless coefficients: 
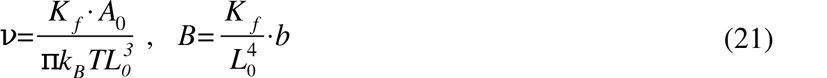

So, the self-consistency equation (19) takes the form below: 
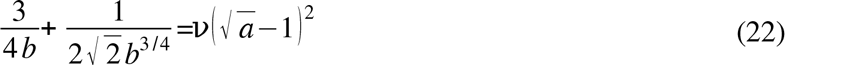

Solving of equation (19) gives the expression for entropic coefficient: 
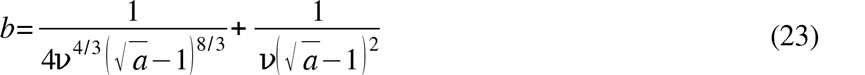

Then, the area per chain *a* in the flat monolayer may be found from equilibrium equation [1,2]. 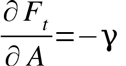 Calculated area per lipid chain and entropic potential coefficient for various lipids are represented in (Supplementary table 1.

Finally, we obtain the expressions for derivatives of entropic coefficient consisted in expression for lateral pressure profile (14).

One can differentiate the selfconsistency equation (19): 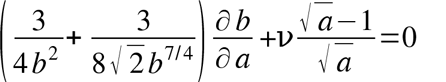 and find expression for the first derivative of entropic coefficient 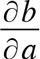:

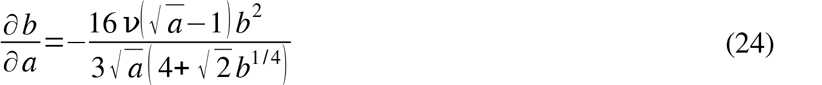

Further, one can redifferentiate obtained equation:
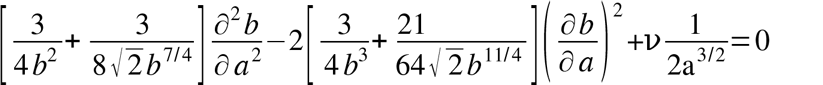 and find expression for the second derivative 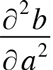:

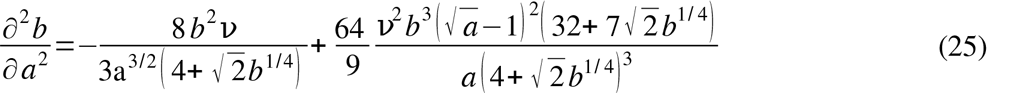

## Results

### Model of bent bilayer membrane

Here we introduce our description of bent bilayer. We consider the pure bending of the membrane (see fig. 2). We assume that the pivotal surfaces are located in the hydrophobic-hydrophilic interfaces of the bilayer membrane [19, 20, 21]. Conservation of volume per lipid causes top layer to become thinner while the bottom one becomes thicker. Below we use notations *A_b_* (*z*) for the area per chain in the inner (‘bottom’) monolayer, and *A_t_* (*z*) for the outer (‘top’) monolayer, respectively (see fig. 2).

As is well known from differential geometry [20], for two parallel curved surfaces at a distance *z*, the area elements are related by the following equation: *dA*’=*dA*·(1-*zJ+z^2^K*), where *J* is mean curvature; *K* is Gaussian curvature. In what follows we disregard Gaussian curvature contribution for the chosen topology of the membrane and write the area change in the form [22]:

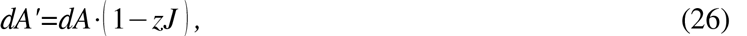

where *J* is the curvature of the surface *dA*.

Using formula (26), one may define the area change via depth coordinate *z* for the ‘bottom’ monolayer in a curved bilayer membrane (the mean curvature of the bottom monolayer is represented by expression: J_b_=−J (1+ *L_b_J*)< 0) and for the ‘top’ monolayer (*J_t_*=*J* (1 – *L_t_J*)> 0): 
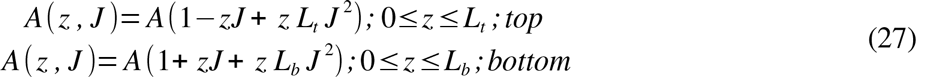

In eq. (27) the origin of *z* is different for ‘bottom’ and ‘top’ monolayer (see fig. 2), *A* – the area per lipid chain in flat monolayer (*J* =0): *A*=*A*_0_·*a*, where *A*_0_ is incompressible area per lipid chain and *a* is dimensionless area per lipid chain.

Using formula (27) we can calculate the derivatives of the area per lipid chain with respect to curvature *J* in each monolayer:

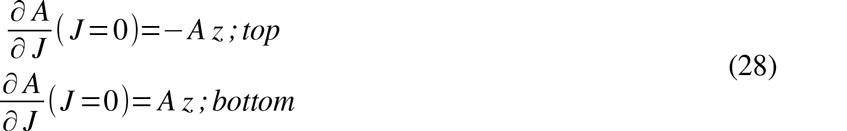

**Figure 2.**
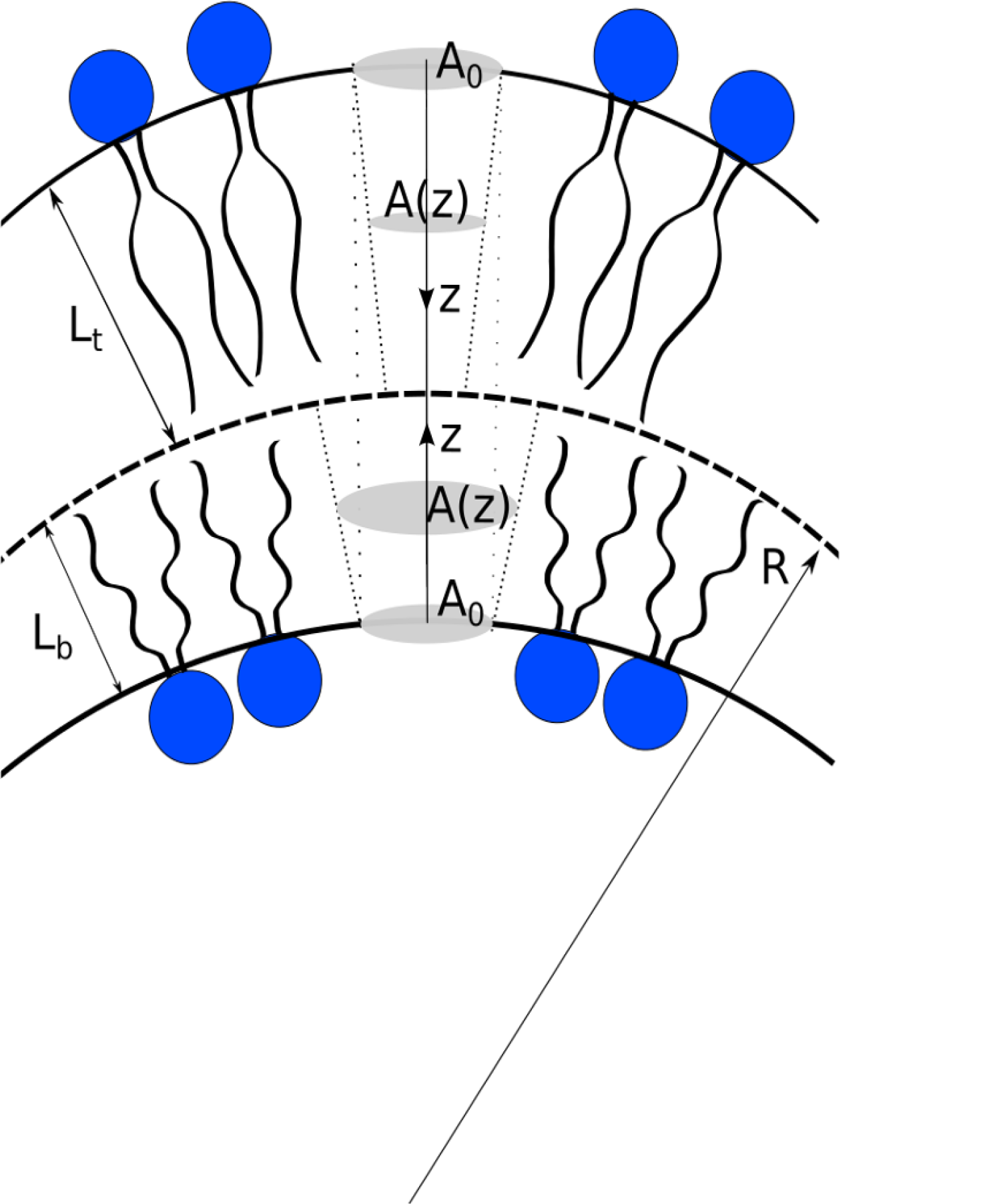
Membrane bending model. *L_b_*–the thickness of the inner monolayer, *L_t_*–the thickness of the outer monolayer; 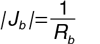 and 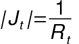–mean curvature of bottom and top cylindrical pivotal surfaces, respectively, where R_b_=R–*L_b_* and *R_t_*–R+L_t_–the curvature radii of these surfaces. For spherical surface the mean curvature for monolayers are given by expressions: 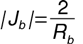, 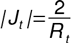. Here *J* is the curvature of the intermonolayer surface. The curvatures of bottom and top pivotal surfaces differ by sign. Under bending area per lipid at the headgroup pivotal surfaces of each monolayer is unchanged with respect to flat membrane provided there is a lipids reservoir or intermonolayer slide.

### The change of lateral pressure profile induced by curvature

In this section we re-derive in the present context the formula (14).

Namely, the perturbation theory can be applied to the conformational energy operator *Ĥ* (3) and the first correction to the lateral pressure profile due curvature can be calculated.

In the case of bent membrane the entropic repulsion coefficient *B* of the flat bilyer in formula (3) acquires the curvature induced depth-dependent correction *B*^[1]^(*z,J*). Thus, the perturbation of ‘Hamiltonian’ is *Ĥ*^[1]^:

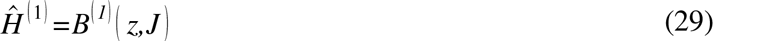

Further function *B*^[1]^(*z*) constituting the perturbation operator in (29) can be obtained in the first approximation by the Taylor’s expansion of *B* in powers of the curvature *J*: 
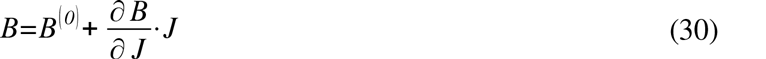

In this case derivative of the entropic potential 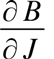 is taken at zero curvature *J*=0.

The first correction to entropic potential is represented by the formula:

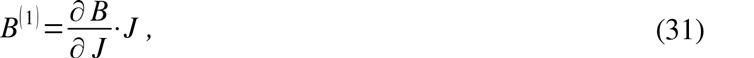

where derivative of *B* with respect to curvature 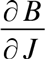 equals:

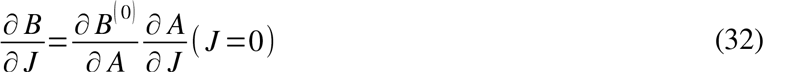

We apply perturbation theory 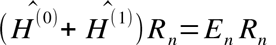 to obtain the first correction 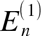 to eigenvalue 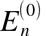 [23]: 
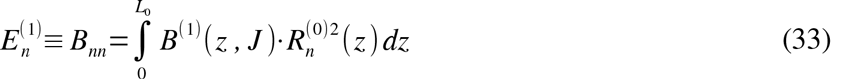

Then we can re-write the free energy for curved membrane in the operator form 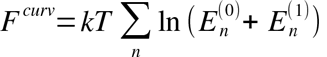 and differentiating it with respect to area per chain *A(z)* we can find lateral pressure profile in curved membrane 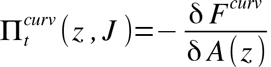: 
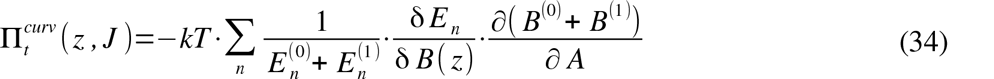

Using formula (33) one can calculate the variational derivative 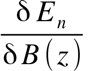 and substituted obtained expression in (34) and then distinguish the lateral pressure profile in flat membrane 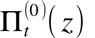 (see eq. (14)) and the correction to the lateral pressure profile due curvature 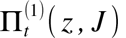: 
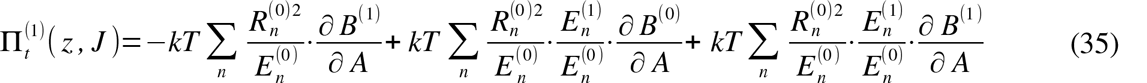

As one can see from equations (31), (32) and (33) the last term in formula (35) is proportional to the second order curvature *J*^2^ and it may be vanished in the first approximation for the case of small bending. So, the final expression to the correction to lateral pressure due curvature is represented by formula: 
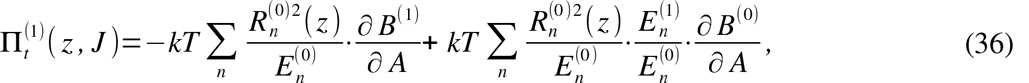
where 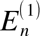 and 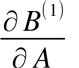 can be re-written as: 
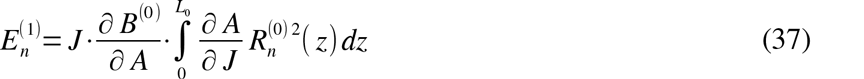

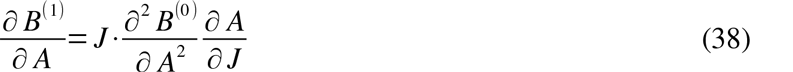

Expressions for derivatives of *B*^[0]^ with respect to *A* may be re-written through dimensionless ones: 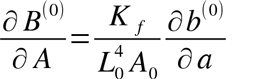 and represented in Supplementary table 1. The corrections to the lateral pressure profile 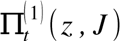 in the inner and the outer monolayers of the bent bilayer can be written in the form: 
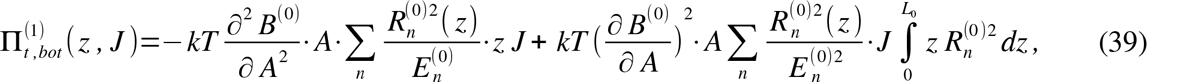
where *A*=*a*·*A*_0_–the area per lipid chain in flat bilayer.

In expression (39) variable *z* varies in the interval *z*∊[0*; L_b_*], where 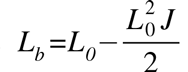 is the thickness of the inner (‘bottom’) monolayer that follows from the volume per lipid conservation (see fig. 2).

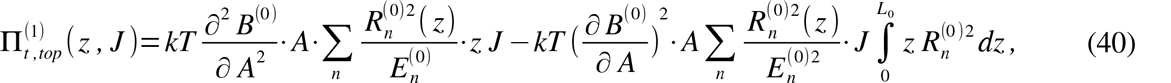

In expression (40) variable *z* varies in the interval: *z*∊[0*; *L*_t_*], where 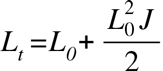 is the thickness for the outer (‘top’) monolayer, derived from conservation condition of the volume per lipid (here *J* is curvature of the intermonolayer surface – see fig. 2)

The results (39)-(40) are plotted in fig. 3.

As one can note the bending shifts position of the peak of lateral pressure profile along the *z*-axis. This is caused by the fact that the top and bottom monolayers acquire different thicknesses due to the bending (see Fig. 2).

Inside the inner layer *z*∊[0; *L_b_*] of the bent bilayer the area per one tail is greater than in the flat membrane (see eq. (27)), therefore, the entropic repulsion is weakened, and, hence, the lateral pressure is lowered (see fig. 3). In the outer layer *z*∊[0*; L_t_*] of bilayer inter-tails space is compressed and entropic repulsion between tails, i.e. lateral pressure, raises up.

Obtained from our microscopic model the lateral pressure profile in hydrophobic part of the bilayer (fig. 3) agrees qualitatively well with results of molecular dynamic simulation (see fig.2a in work [24]) and with our previous obtained calculated pressure profile [25].
Also our asymmetrical lateral pressure profile is well consistent with earlier obtained computational calculations by authors [26] obtaining the lateral pressure profile imbalance in the DOPC bilayer with the asymmetric addition of non-bilayer lipids into one of the monolayers. Also authors of this article [26] and other work [6, 27] investigated that asymmetric incorporation of conical lipids into bilayer was shown to activate mechanosensitive channels (particularly, the simplest model system such a mechanosensitive channels of large conductance) in the absence of external surface tension. In next section we’ll check analytically this statement.

**Figure 3.**
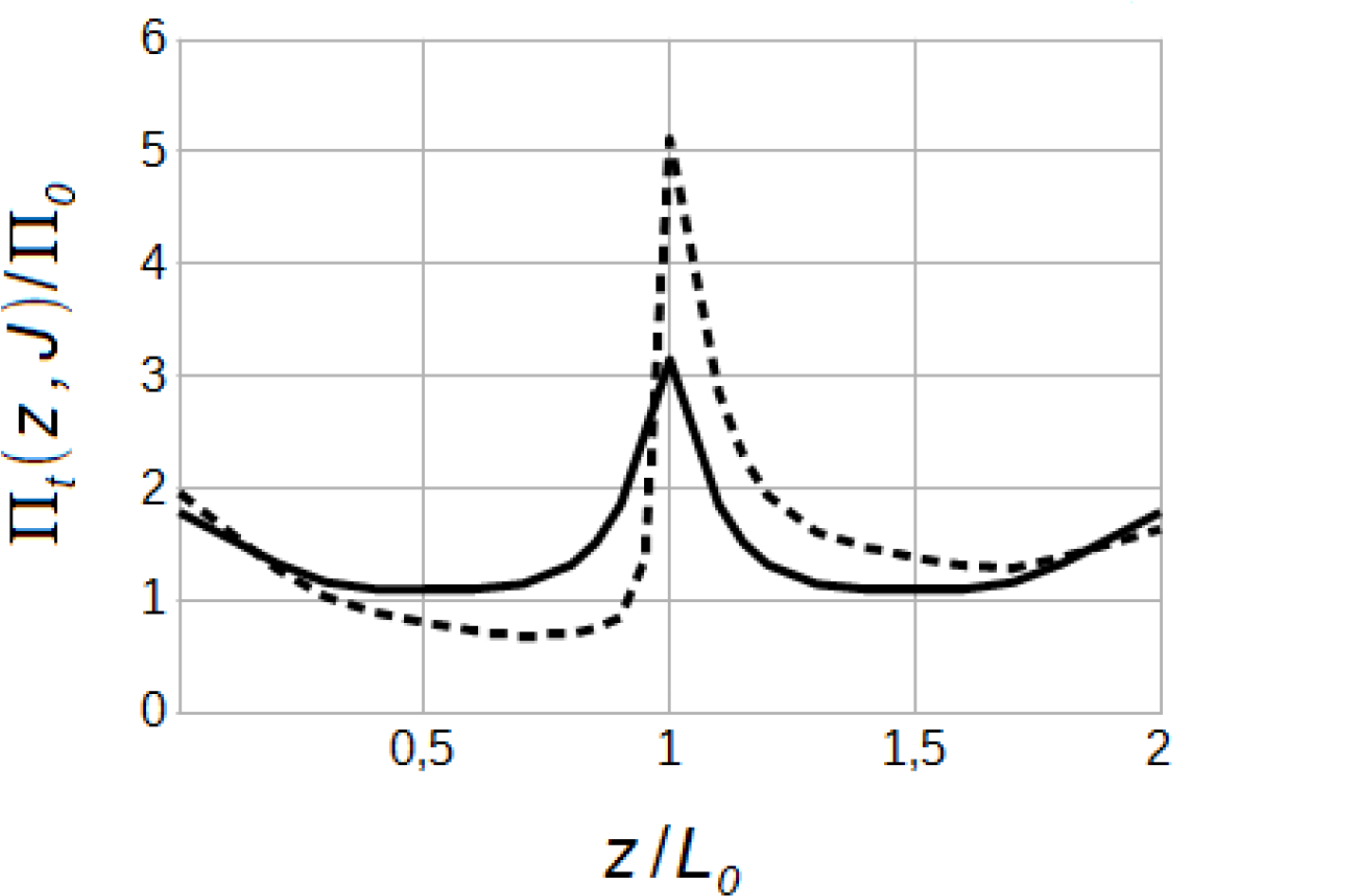
Lateral pressure profiles in hydrophobic part of DPPC bilayer, normalized with П_0_ =δ*/L*_0_. Lateral pressure in the flat membrane (solid line), lateral pressure profile in lipid bilayer with finite curvature 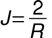 (dotted line), where *R*=20*nm* – the fixed curvature radius. Here γ=50*dyn/cm* is inter-facial surface tension, *L*_0_=1,5*nm* - the monolayer thickness.

### Mechanosensitive channel gating

The conformation shape of MscL was found to change between cylindrical (closed) and conical (open) shapes [28]. Other authors [29] assumed that conical shape of channel corresponds to its intermediate state, but hour-glass shape corresponds to its open state (see fig. 4). To estimate the effect of pressure profile on membrane proteins, we followed the approach introduced in articles [4], [13], [30] where was shown that lateral pressure profile affects the functioning of the proteins embedded in lipid membrane and was calculated numerically the work on MscL opening: 
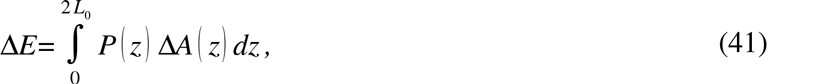
where *P*(*z*) – lateral pressure in hydrophobic part of the bilayer, ΔA (*z*) — the changing of the channel area during its opening process. The mechanosensitive channel opening of pore was experimentally determined [12] to have a diameter of *d* =2,5 *nm*.

**Figure 4.**
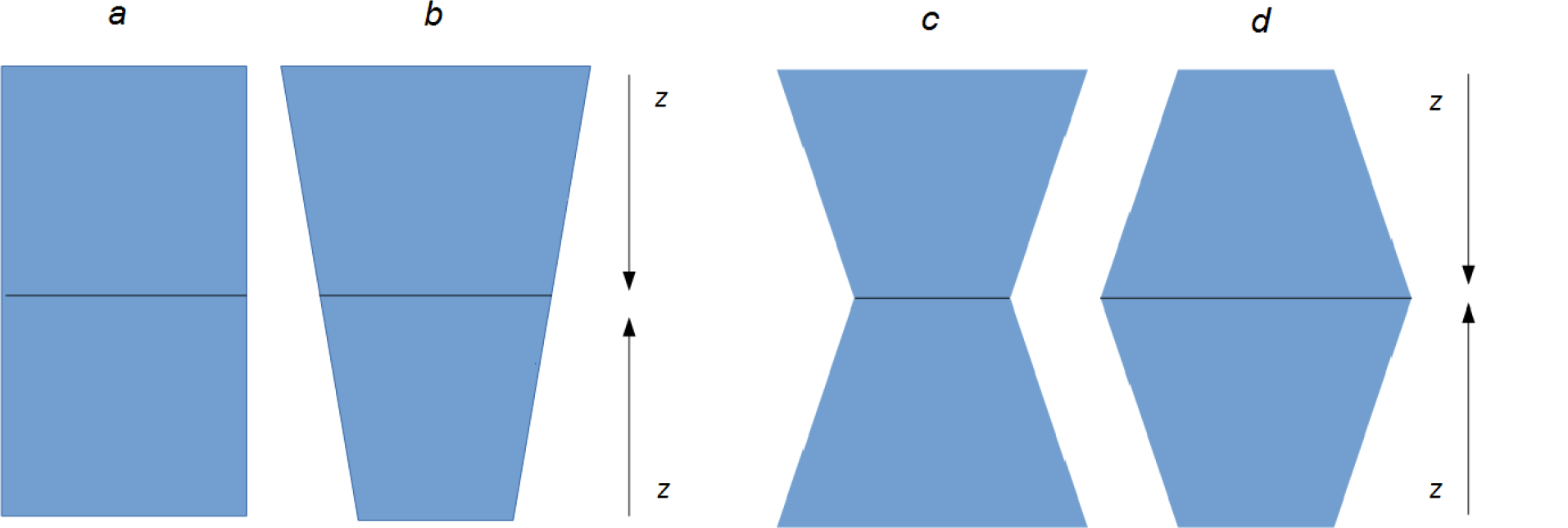
The modeling of MSC shape. *a*: the cylindrical channel; *b*: conical shape of the channel; *c, d* the hour-glass shape of the channel; the cross-sectional area in the middle of the mechanosensitive channel doesn’t change.

The authors of work [5] using the molecular dynamic methods have shown that lipid bilayer bending is the cause why MSCs can to be opened. Our aim in this section is to test analytically this hypothesis and continue our previously started calculation [31] and obtain the expression for energy difference between two various states of the mechanosensitive channel.

At the beginning we define the area difference between states of MSC using experimental data [12]. The cross-sectional area of channel *A*(*z*) is decomposed into two components: a constant mid-plane area *A_m_* and z-dependent shape variation Δ *A* (*z*) around *A_m_*, and for conical shape of the channel we obtain asymmetric area changing with respect to middle of the bilayer, but for hour-glass one – symmetrical area changing [29].

Then for the conical shape of the mechanosensitive channel (see fig. 4b) we obtain the next area changing: 
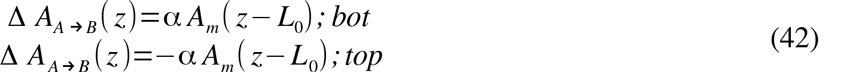
also for hour-glass conformation of the channel (see fig. 4c): 
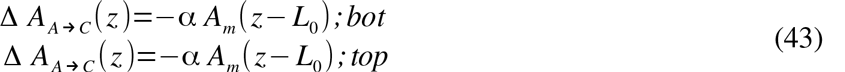
and for hour-glass conformation of the channel (see fig. 4d), respectively: 
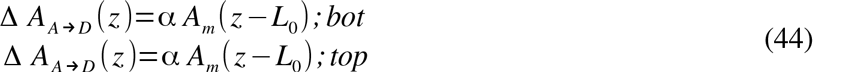

In eq. (42)–(44) zero of axis *z* is in the pivotal surfaces (see fig. 1, 2), 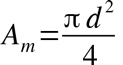 is changing of MSC area in the middle of bilayer and the slope is [12]: 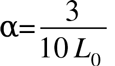, where *L*_0_ is the monolayer thickness.

At last, we can write the energy difference between two various states of channel in the following form [32]: 
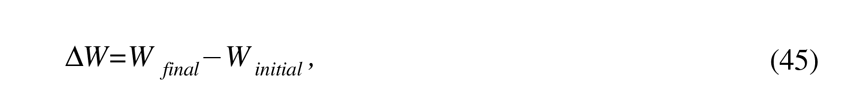
where *W _final_* and *W_initial_* – the energy of final and initial state of lipid environment and the channel.

The comparison of obtained results for the energy difference between two states of mechanosensitive channel in conical and hour-glass models are represented in table 1. Decreased opening barrier due to the lateral pressure imbalance and the spontaneous bending are shown separately in Supplementary figures 1, 2. The figure S1 shows that when MSC changed its shape from cylindrical to conical one, the bending energy of bilayer decreases by 5 *k_B_T* average. During this process the stretching energy is increasing by amount about 10 *k_B_T* (see tables 1, 2). These estimate agrees qualitatively well with results of molecular dynamics simulations presented in [4, 32, 33] where values of the energy difference between open and closed MSC states is about 0-10*k_B_T*. The figure S2 shows that when the channel has changed its shape from cylindrical to hour-glass one, the spontaneous bending energy decreases by amount 8-15 *k_B_T* while the membrane hydrophobic energy increases by amount about 15-30*k_B_T* (see table 2).

Analytical calculation of the bending modulus and spontaneous bending moment for the case of small bending are represented in Supplementary text 1. The obtained results are good agreement with previously obtained result for spontaneous bending moment *κ_m_·J_s_=(*-151 ±6)×10^-13^*J/m* [24] and bending modulus of lipid monolayer [24], [34] (see table 3).

The analysis of the relation between bending and stretching moduli of the bilayer is represented in table 4. Usually experimental value for bending modulus for different lipid bilayers vary between (0,1-6)×10*^-19^J* [35–37]. The relation between bilayer bending modulus and area stretching modulus of the bilayer obtained from our calculation is expressed by the formula: 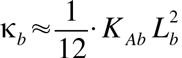, where *L_b_*=2 *L*_0_ – the bilayer thickness and κ_*b*_ = 2 κ_*m*_ the bending modulus for the bilayer. This condition is considered as correct one for symmetrical monolayers [37, 38].

**Table 1.**
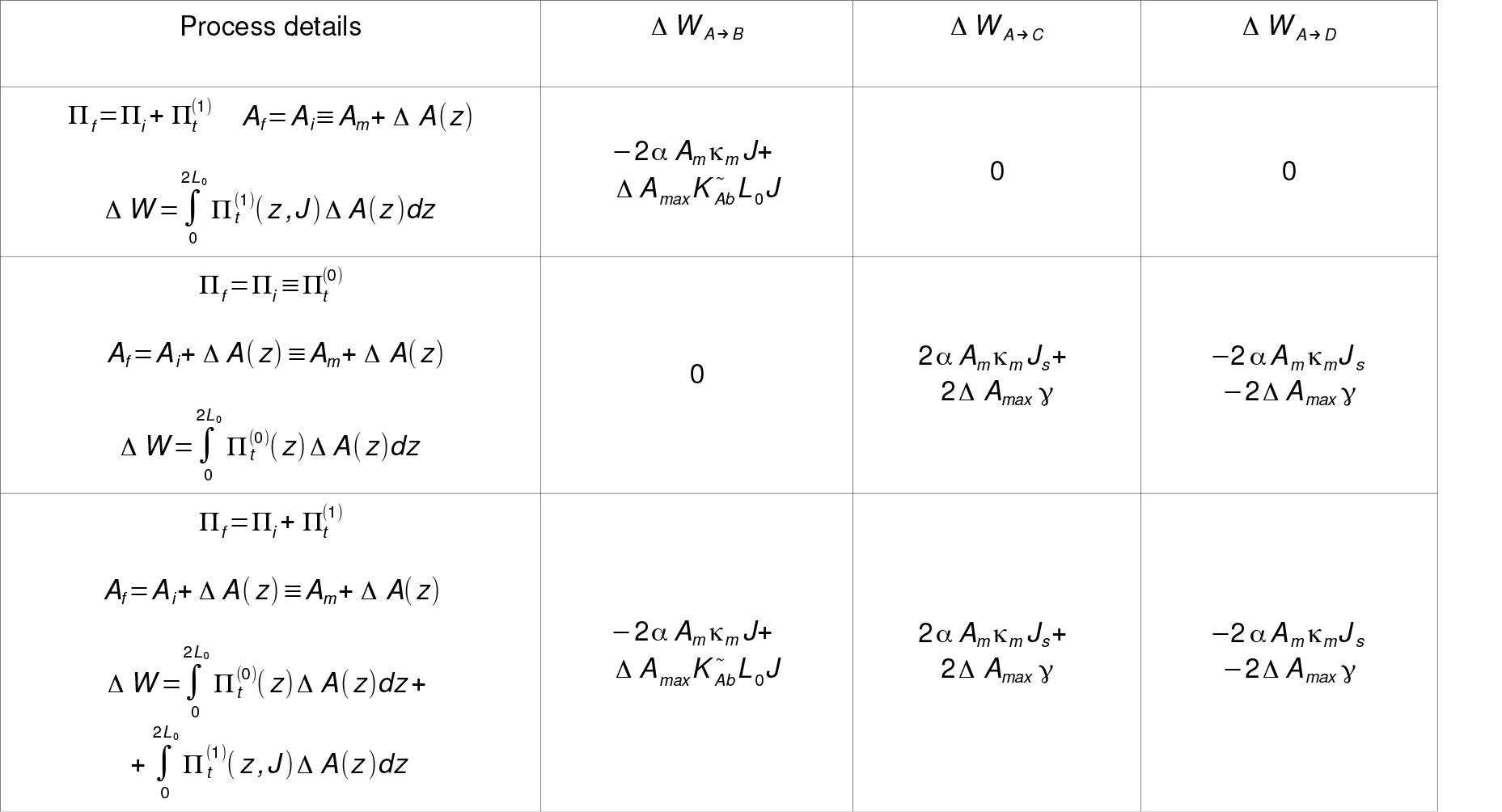
The energy difference between two states of the mechanosensitive channel in conical and hour-glass models. Here K*_m_* – the bending modulus of the monolayer and K*_m_*J*_s_* - the first spontaneous bending moment (see Supplementary text 1), *L*_0_ – the monolayer thickness, *J* - the curvature of intermonolayer surface (see fig. 2), γ - the inter-facial surface tension. *A_m_=*4,9*nm^2^* is the area of MSC in the middle of bilayer. The last line in this table shows the mixed process in which at first step the spontaneous curvature was induced in one of the monolayers of bilayer and then this bilayer was bent in addition. Index *i* means the initial state, and *f* means the final state of the mechanosensitive channel and its environment (lipid bilayer). The channel area changing at the inter-facial region is 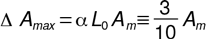 (see eq. (42)–(44)). 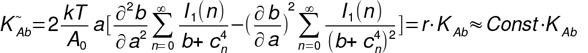 – re-normalized area stretching modulus of the bilayer, where expression for the integral *l*_1_ (*n*) one can find in Supplementary text 1.

**Table 2.** The calculations of the energy difference between two conformations of MSC in DPPC bilayer environment. (*A*_0_=0,2 *nm*^2^ is an incompressible area per lipid chain, *T_0_=300K* is temperature, *L*_0_=1,5*nm* is the monolayer thickness). The result for hour-glass model, fig. 4d is different sign in comparison with hour-glass, fig. 4c (see table 1). For calculation the first bending moment of the lateral pressure was taken the curvature radius *R=20nm* and curvature for spherical surface. 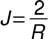. Here γ is the inter-facial surface tension. As one can note that transitions between channel shapes are characterized by relation between energy differences: *Δ W_A→B →C_*=*Δ W_A→B_*+ Δ *W_B→C_*≡Δ *W_A→C_*.

| Contribution into energy difference | $\Delta W_{A \rightarrow B}, k_B T$ | | | $\Delta W_{A \rightarrow C}, k_B T$ | | | $\Delta W_{B \rightarrow C}, k_B T$ | | |
| --- | --- | --- | --- | --- | --- | --- | --- | --- | --- |
| $\gamma, \text{ mN/m}$ | 30 | 40 | 50 | 30 | 40 | 50 | 30 | 40 | 50 |
| $\vert 2\alpha A_m \kappa_m J_s\vert $ | 0 | 0 | 0 | -8,15 | -10,85 | -13,56 | -8,15 | -10,85 | -13,56 |
| $\vert 2\Delta A_{max} \gamma\vert $ | 0 | 0 | 0 | 21,33 | 28,44 | 35,55 | 21,33 | 28,44 | 35,55 |
| $\vert 2\alpha A_m \kappa_m J\vert $ | -2,08 | -3,02 | -4,04 | 0 | 0 | 0 | 2,08 | 3,02 | 4,04 |
| $\vert \Delta A_{max} K_{Ab} L_0 J\vert $ | 2,42 | 3,52 | 4,72 | 0 | 0 | 0 | -2,42 | -3,52 | -4,72 |
| $\Delta W_{\Sigma}$ | 0,34 | 0,5 | 0,68 | 13,18 | 17,59 | 21,99 | 12,84 | 17,09 | 21,31 |

**Table 3.** The spontaneous bending moment κ*_m_ J_s_* and the bending modulus for the lipid monolayer K*_m_* calculated analytically within the microscopic model of flexible strings. The input parameters of the lipid chain are: *A*_0_=0,2 *nm*^2^-incompressible area, *T*_0_=300*K* the temperature, γ the surface tension at the interface between the polar and hydrophobic regions of the membrane. Here *L*_0_=0,9*nm* the monolayer thickness of DDPC(10:0) lipids, *L*_0_=1,1*nm* the monolayer thickness of DLPC(12:0) lipids and so on.

| Lipid | $\gamma, mN/m$ | $\kappa_m \times 10^{-20} J$ | $\kappa_m J_s \times 10^{-12} J/m$ |
| --- | --- | --- | --- |
| DDPC (10:0) | 30 | 1,5743 | -10,5144 |
|  | 40 | 2,0631 | -12,8678 |
|  | 50 | 3,0956 | -17,2432 |
| DLPC (12:0) | 30 | 2,3397 | -12,7126 |
|  | 40 | 3,423 | -16,8399 |
|  | 50 | 4,6281 | -20,9930 |
| DMPC (14:0) | 30 | 3,2790 | -14,9388 |
|  | 40 | 4,7141 | -19,6062 |
|  | 50 | 6,4492 | -24,7940 |
| DPPC (16:0) | 30 | 4,383 | -17,1893 |
|  | 40 | 6,376 | -22,8856 |
|  | 50 | 8,5235 | -28,5995 |

Finally we can point out that our analytical calculation allows estimate quantitatively how changes in the lateral pressure of the bilayer influences on MSC gating mechanism. Altering in pressure profile produces the changing in the locking energy barrier of MSC 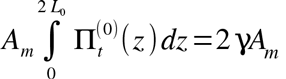 (see eq. (11)-(12)). And of course, if changes in lipid composition induces spontaneous curvature, then these changes would be produce a significant conformational energy change.

**Table 4.**
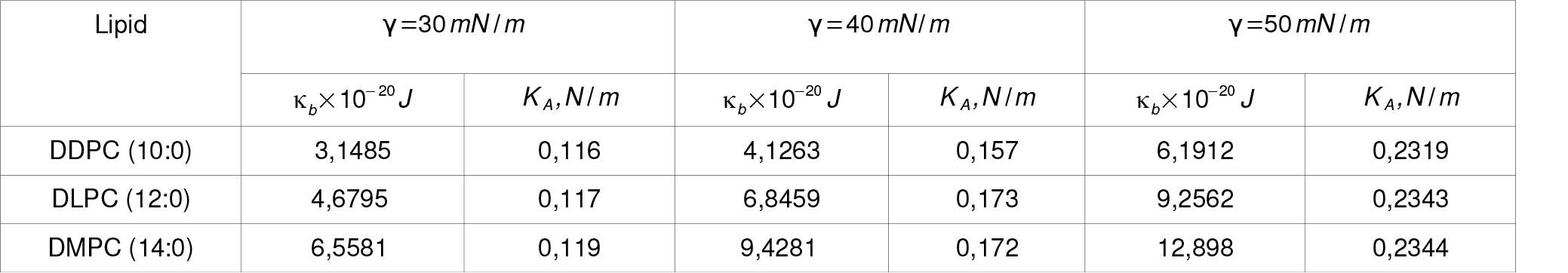
The calculation of the bending modulus for various bilayer and its comparison with stretching modulus in the flexible strings model. κ*_b_* – the bending modulus of the bilayer, γ – the inter-facial surface tension, 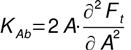 – the stretching modulus of the bilayer. Expression for stretching modulus in flexible strings models is: 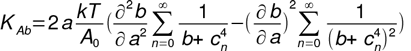. Here *L*_0_=0,9*nm* – the monolayer thickness of DDPC(10:0) lipids, *L*_0_= 1,1 *nm* – the monolayer thickness of DLPC(12:0) lipids and so on.

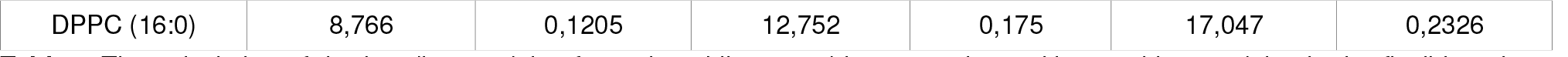

Also, one can note that the great inter-facial tension γ corresponds to a greater change in the energy barrier because the bending modulus K*_m_* and the entropic potential *b* grow up with increasing γ. The energy difference for various lipids is represented in Supplementary table 2. As one can see from table S2, MSCs have been shown to be sensitive to inter-facial surface tension and spontaneous bending moment values. These results are well consistent with numerical calculations of previous study [4, 32]. The table S2 shows that for fixed slope of the channel 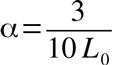 we obtain that the absolute value of work associated with embedding of MSC in its hour-glass configuration into the flat membrane does not depend on the monolayer thickness and depends majorly on the re-normalized inter-facial surface tension Δ *W*_A↔C_≈βΔ *A_max_·γ*, where β=1,235±0,005 – the renormalization constant. This is caused by the fact that spontaneous bending energy 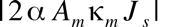 (see eq. (10), (12), (S8)) lowers/increases the hydrophobic energy 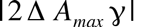 to the permanent value. This result is well consistent with previous calculation [32]. Of course, one may conclude that bending of membrane associated with simple cylinder-cone-like shape channel transition doesn’t produce a great changing in energy difference (see tables 2, S2). Moreover the embedding of MSC in the bent membrane consisting of lipids with short hydrocarbon chains corresponds probably to the intermediate state of the channel, but embedding in long-tails lipid environment corresponds most likely to the closed state of the channel (table S2). This is caused by the fact that for the channel a probability of being open 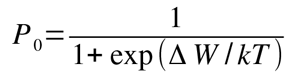 is reducing with increasing the energy difference (Δ *W* is the energy difference between open and closed states of the channel [11, 32, 39]). At contrast the decreasing of the energy barrier corresponds to the open state of MSC (the negative values of the energy difference corresponds to the opening of the mid-plane pore in the channel).

## Author contributions

A.A.D. did the analytical derivation of all thermodynamic characteristics reported in this article and wrote initial version of the paper. S.I.M. supervised through out the research.

---

**Figure 1.**
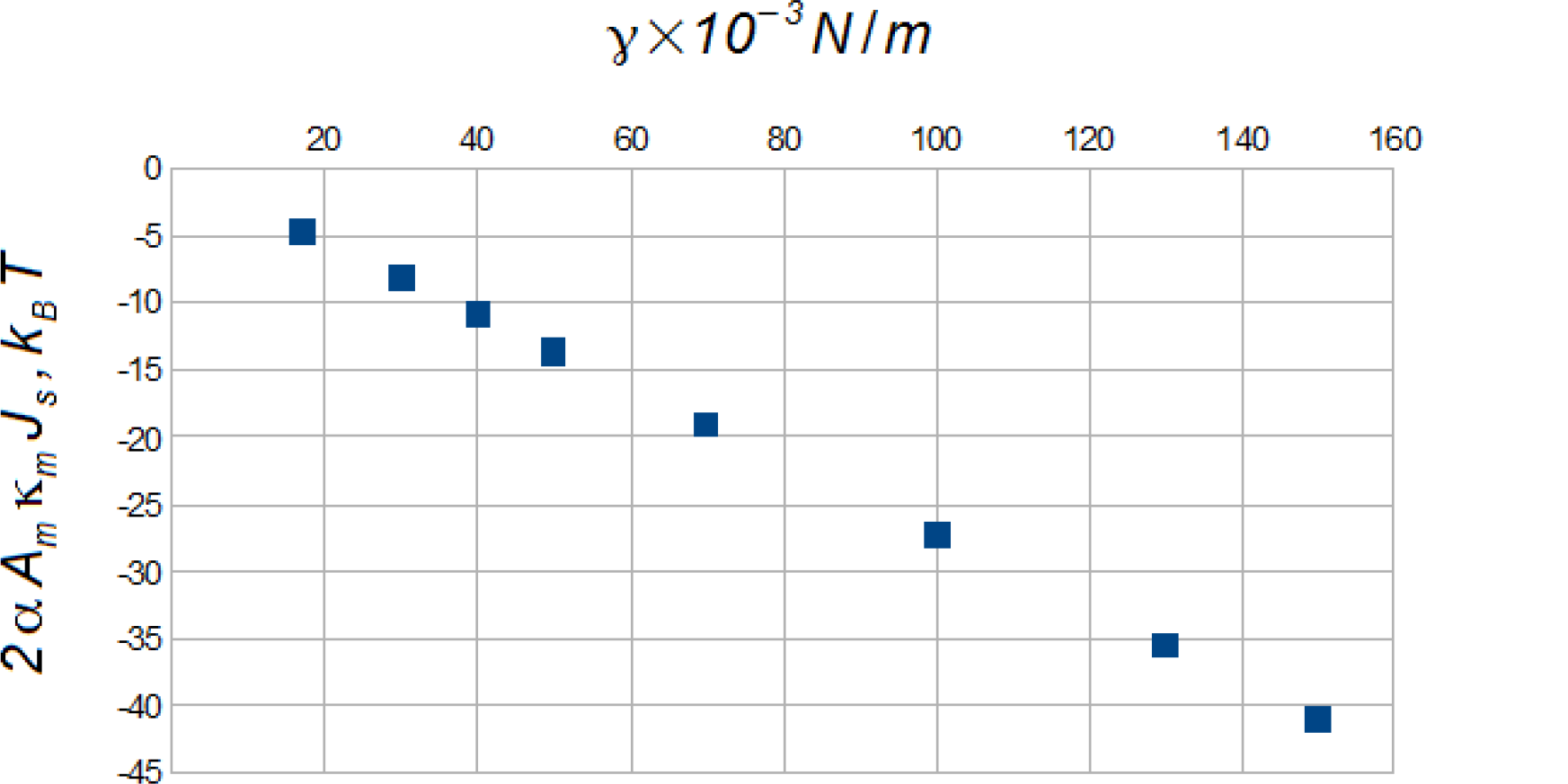
Decreasing opening barrier due to the spontaneous bending of the lipid membrane, at temperature *T*=300 *K*. Here γ - the surface tension between polar and hydrophobic regions and κ*_m_J_s_* – the first spontaneous bending moment of the lateral pressure (see Supplementary text 1), 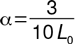 – the slope of MSC, *L*_0_ - the monolayer thickness, *A_m_* - the area of the channel in the middle of bilayer.

**Figure S2.**
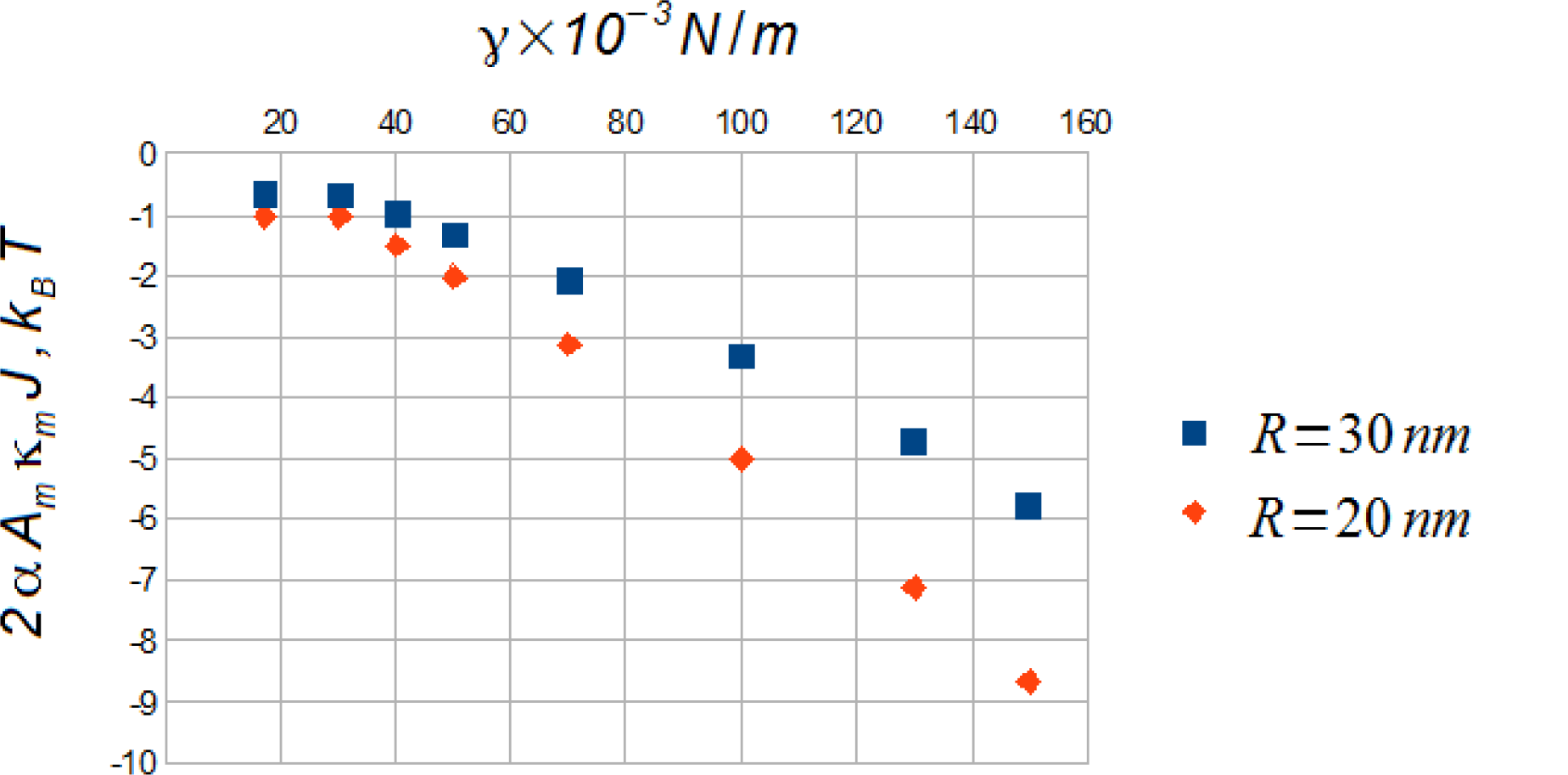
The reduce of the energy difference between two conformational states of MSC embedded in a bent DPPC bilayer. Here γ – the surface tension between polar and hydrophobic regions and *R* – curvature radii, κ*_m_J* - the first bending moment of the lateral pressure (see Supplementary text 1). Here was used the curvature for cylindrical surface 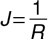 (see fig. 2).

**Table S1.** The parameters of the lipid chain calculated within the flexible string model at temperature *T*_0_=300*K*. *a* - area per lipid chain, *b* – the entropic coefficient, 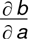 – the first derivative of the entropic coefficient, 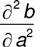 – the second derivative of the entropic coefficient. The fixed parameters for each lipid are: *A*_0_=0,2*nm^2^* – the incompressible area per chain, 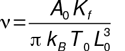– dimensionless parameter and *K_f_=k_B_T_0_L*_0_/3 – the bending rigidity of the lipid chain. We vary *L*_0_ – the monolayer thickness and γ – the inter-facial surface tension. Here *L*_0_=0,9*nm* – the monolayer thickness of DDPC(10:0) lipids, L_0_ = 1,1 *nm* – the monolayer thickness of DLPC(12:0) lipids and so on.

| Lipid parameter | $\gamma, mN/m$ | DDPC (10:0) | DLPC (12:0) | DMPC (14:0) | DPPC (16:0) | DSPC (18:0) | DAPC (20:0) | DBPC (22:0) |
| --- | --- | --- | --- | --- | --- | --- | --- | --- |
| a | 30 | 3,436 | 3,544 | 3,653 | 3,76 | 3,864 | 3,963 | 4,057 |
|  | 40 | 3,055 | 3,071 | 3,189 | 3,265 | 3,355 | 3,439 | 3,518 |
|  | 50 | 2,678 | 2,774 | 2,864 | 2,949 | 3,027 | 3,1 | 3,168 |
| b | 30 | 101,378 | 149,766 | 205,642 | 268,591 | 338,286 | 414,896 | 498,420 |
|  | 40 | 137,974 | 217,865 | 291,904 | 385,138 | 483,27 | 591,465 | 709,560 |
|  | 50 | 201,427 | 291,234 | 394,498 | 510,435 | 640,058 | 782,722 | 938,844 |
| $\frac{\partial b}{\partial a}$ | 30 | -77,547 | -107,912 | -141,283 | -177,231 | -215,328 | -255,640 | -298,035 |
|  | 40 | -126,525 | -197,833 | -250,418 | -320,315 | -387,188 | -458,015 | -532,370 |
|  | 50 | -231,379 | -316,262 | -408,578 | -506,528 | -611,324 | -721,582 | -837,720 |
| $\frac{\partial^2 b}{\partial a^2}$ | 30 | 94,555 | 125,964 | 159,104 | 193,075 | 227,125 | 261,354 | 295,625 |
|  | 40 | 185,44 | 291,313 | 351,076 | 436,555 | 508,486 | 580,678 | 652,765 |
|  | 50 | 425,664 | 555,477 | 686,855 | 815,909 | 946,176 | 1075,656 | 1206,223 |

**Table S2.** The calculated energy difference between two conformational shape of the mechanosensitive channel for different transition and various lipids environment. Here Δ *W_A→B_* – the energy difference for channel transition from cylindrical shape *A* to conical shape *B* and other energy differences may be defined similarly (see fig. 4) and γ – the inter-facial surface tension. One can note that transitions between channel shapes are characterized by relations: Δ *W_A→B→C_=Δ W_A→B_* + Δ *W_B→C_≡Δ W_A→C_*, Δ *W_A→D_=-Δ W_A→_*,Δ *W_A→B→D_=Δ W_A→B→_+Δ W_B→D_=-Δ W_A→B→C_=-Δ W_A→C_* (see table 1). For the calculation the first bending moment was taken the curvature radius R=20*nm* and curvature for spherical surface 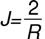. Here *L*_0_=0,9*nm* – the monolayer thickness of DDPC(10:0) lipids, *L*_0_=1,1 *nm* – the monolayer thickness of DLPC(12:0) lipids and so on.

| $\Delta W, k_B T$ | $\gamma, mN/m$ | Lipid | | | | | | |
| --- | --- | --- | --- | --- | --- | --- | --- | --- |
|  |  | DDPC (10:0) | DLPC (12:0) | DMPC (14:0) | DPPC (16:0) | DSPC (18:0) | DAPC (20:0) | DBPC (22:0) |
| $\Delta W_{A \rightarrow B}$ | 30 | 0,17 | 0,22 | 0,27 | 0,34 | 0,4 | 0,46 | 0,52 |
|  | 40 | 0,22 | 0,32 | 0,41 | 0,5 | 0,59 | 0,67 | 0,75 |
|  | 50 | 0,34 | 0,45 | 0,57 | 0,68 | 0,78 | 0,88 | 0,98 |
| $\Delta W_{A \rightarrow C}$ | 30 | 13,02 | 13,11 | 13,16 | 13,18 | 13,2 | 13,2 | 13,2 |
|  | 40 | 18,09 | 17,56 | 17,72 | 17,59 | 17,6 | 17,59 | 17,58 |
|  | 50 | 21,93 | 21,98 | 21,99 | 22 | 21,98 | 21,97 | 21,95 |
| $\Delta W_{B \rightarrow C}$ | 30 | 12,86 | 12,9 | 12,89 | 12,85 | 12,8 | 12,74 | 12,67 |
|  | 40 | 17,87 | 17,23 | 17,31 | 17,1 | 17,01 | 16,92 | 16,84 |
|  | 50 | 21,59 | 21,53 | 21,42 | 21,32 | 21,2 | 21,09 | 20,98 |
| $\Delta W_{B \rightarrow D}$ | 30 | -13,19 | -13,33 | -13,43 | -13,52 | -13,6 | -13,66 | -13,72 |
|  | 40 | -18,32 | -17,88 | -18,12 | -18,09 | -18,18 | -18,26 | -18,33 |
|  | 50 | -22,26 | -22,43 | -22,56 | -22,67 | -22,76 | -22,85 | -22,93 |

## Supplementary text 1

The bending modulus are related to the first moment of the lateral pressure profile in bent monolayer [3, 34]: 
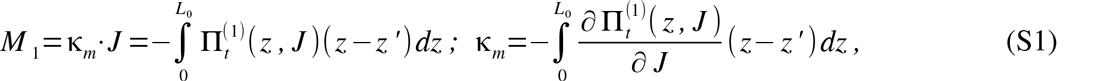
where *L*_0_ is the monolayer thickness and *z’* indicates the pivotal surface position.

The so named first spontaneous bending moment is given in the following form [34]:

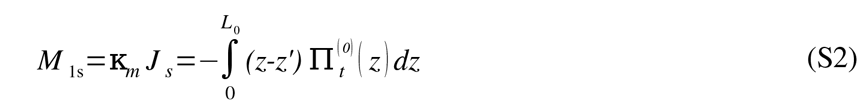

At the beginning we consider one monolayer of the lipid membrane, for example the ‘bottom’ (inner) monolayer (*z*’=0), and calculate analytically the first lateral pressure moment and bending modulus (S1). Substituting formula for the correction to lateral pressure profile for inner monolayer (39) into eq. (S1) and integrating by *z* one can obtain the following expression in dimensionless form (see eq. (21)): 
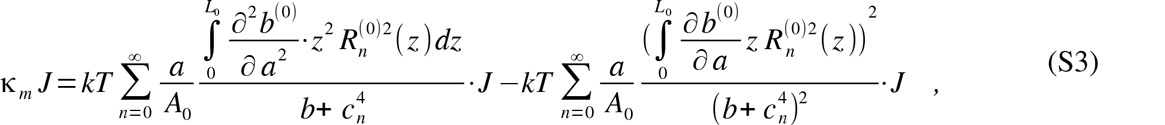
where *A* = *A_0_·a* – the area per lipid chain in a flat membrane.

Computing integrals in (S3) one can obtain the following analytic expression for bending modulus of monolayer: 
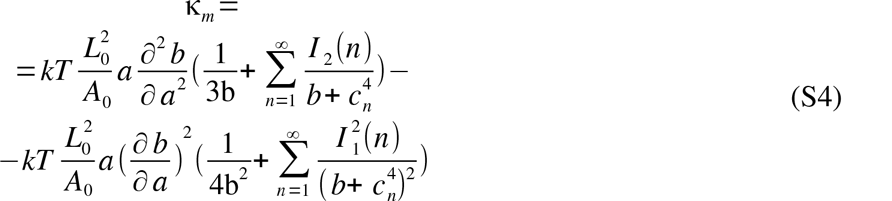

Here expressions for the integrals are represented in the form (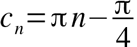): 
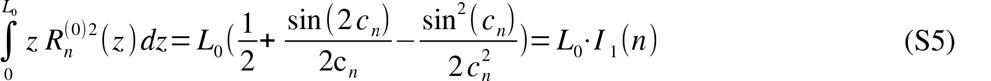

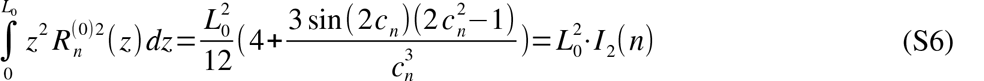

Substitutung expression for lateral pressure in the flat bilayer (14) into (S2) one can calculate the spontaneous bending moment for inner monolayer K_m_ · *J*_s_: 
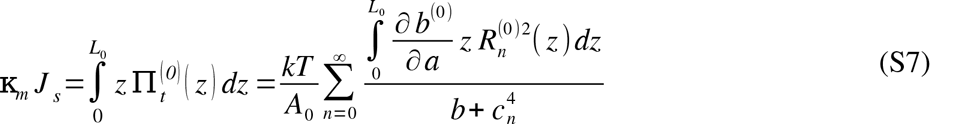

And after the substituting (S5) into (S7) one can obtain the analytic formula for the spontaneous bending moment: 
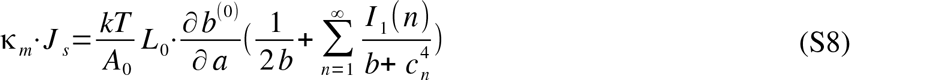

